# Detecting gene subnetworks under selection in biological pathways

**DOI:** 10.1101/128306

**Authors:** Alexandre Gouy, Joséphine T. Daub, Laurent Excoffier

## Abstract

Advances in high throughput sequencing technologies have created a gap between data production and functional data analysis. Indeed, phenotypes result from interactions between numerous genes, but traditional methods treat loci independently, missing important knowledge brought by network-level emerging properties. Therefore, evidencing selection acting on multiple genes affecting the evolution of complex traits remains challenging. In this context, gene network analysis provides a powerful framework to study the evolution of adaptive traits and facilitates the interpretation of genome-wide data. To tackle this problem, we developed a method to analyse gene networks that is suitable to evidence polygenic selection. The general idea is to search biological pathways for subnetworks of genes that directly interact with each other and that present unusual evolutionary features. Subnetwork search is a typical combinatorial optimization problem that we solve using a simulated annealing approach. We have applied our methodology to find signals of adaptation to high-altitude in human populations. We show that this adaptation has a clear polygenic basis and is influenced by many genetic components. Our approach improves on classical tests for selection based on single genes by identifying both new candidate genes and new biological processes involved in adaptation to altitude.

## INTRODUCTION

Understanding the genetic basis of adaptation remains a central theme of evolutionary biology. Adaptation is typically viewed as involving selective sweeps that drive beneficial alleles from low to high frequencies in a population, lowering genetic diversity and increasing linkage disequilibrium near the selected region (1-3). Numerous statistical tests have been developed to detect selection from genomic data based on a simple selective sweep model (reviewed in De Mita et al. (4)). Therefore, most work in humans and other species has focused on identifying signals of strong selection at individual loci (5). These methods have been quite successful in humans to identify loci involved in several adaptations such as diet, altitude, disease resistance, and pigmentation (reviewed in Vitti et al. (6)). However, examples of adaptation due to a selective sweep at a single locus remain relatively rare in human populations. Therefore, some authors have argued that adaptation events could occur by the evolution of polygenic traits rather than via the fixation of single beneficial mutations (7-9).

Recent Genome-Wide Association Studies (GWAS) in various model organisms have confirmed that variation at many important traits is controlled by a large number of loci scattered throughout the genome, e.g. human height (10,11). Selection acting additively on this kind of traits could therefore lead to small shifts in allele frequencies (8). This verbal model has been studied analytically, showing that in some cases, polygenic selection may indeed lead to subtle shifts in allele frequencies (12-14). However, these small allele frequency changes may remain below the detection limit of most of outlier detection methods (15). Therefore, the generality of conclusions drawn from significant tests can be seriously challenged because phenotypic traits exhibiting clear-cut molecular signatures of selection may represent a biased subset of all adaptive traits (16). Another caveat of classical genome scans for selection is that lists of candidate genes are sometimes difficult to connect to a particular adaptive mechanism, since SNP-level results are unlikely to reveal complex mechanisms of adaptation, given the lack of signal of small-effect alleles. It seems therefore necessary to consider alternative approaches to study the genetic basis of adaptation of complex traits.

Current approaches to detect selection acting on polygenic traits rely mostly on quantitative genetics models. Classical quantitative genetics approaches are not based on genetic data, but on an explicit description of continuous phenotypes (e.g. height, body mass index, fertility, etc.). These methods have strong theoretical foundations, and allow one to disentangle the genetic from the environmental variance by taking into account the heritability of the traits, and therefore to detect shifts in the distribution of the phenotype under selection (17). But these methods do not permit to identify the genetic basis of adaptation, and other approaches must be considered to associate genetic data to quantitative traits responding to selection. Correlative approaches have emerged where associations between a genotype and various environmental variables are tested (e.g. (18-21)). Finally, recent approaches have tried to estimate selection coefficients from GWAS data (9,22,23), but all these methods need some phenotypic measures of the tested individuals or associated environmental data, which can be sometimes difficult to obtain.

In contrast to a gene-centric approach, some studies have considered testing if a set of genes as a whole is yielding signals of selection (7,24). Indeed, different genes within pathways (i.e. molecular networks leading to a given biological function) may interact to produce a given phenotype (25,26), and therefore be under the same selective pressure. Finding sets of outlier interacting genes can be achieved using gene-set enrichment methods (e.g., (27,28)). The idea is to assign a score (i.e. proxy for selection) to each gene within a biological pathway (i.e. gene-set) and to test if the distribution of scores within the pathway is significantly shifted towards extreme values (7). This approach has successfully identified candidate pathways involved in various human adaptations, such as response to pathogens (7), or adaptation to altitude (24). However, this gene-set enrichment approach mainly identifies pathways where all its members show a shift in the distribution of a given tested statistic. It might thus be underpowered to find more subtle signals, where only a subset of genes is under selection in a large pathway, which is a more likely situation than assuming that all the genes in a pathway have responded to selection.

To address this problem, network analysis can provide new insights into the genetics basis of adaptation. In the last few years, network-based approaches have spread into a large number of research areas, and were successfully used to solve a wide range of biological problems; e.g. gene expression studies (29,30), GWAS (26) or evolutionary biology (31,32). Here, we present a new network-based method to detect polygenic selection in natural populations. The general idea is to search for subnetworks of interacting genes within biological pathways that present unusual features. This search is a typical combinatorial optimization problem that can be solved using a heuristic approach like simulated annealing (29,33). We implemented such an algorithm to search for high-scoring subnetworks of genes in biological pathways, and we developed a testing procedure that explicitly takes into account the optimization process involved in this search. After studying the sensitivity and precision of our method with simulated data, we reanalysed data from a previous study looking for convergent adaptation to altitude in Tibetans an Andeans (24). As compared to the original study, we discover new genes and biochemical functions potentially related to adaptation in these human populations. Our method can thus complement classical genome scans by providing functional information and discovering new genes with weaker effects that are involved in complex selective processes. Finally, we discuss the limits and potential improvements, as well as other possible applications of our methodology.

## MATERIAL AND METHODS

### Pathway databases and conversion to gene networks

We considered biological pathways as gene networks. More formally, we define a gene network as a graph *G*(*V*,*E*), where *V* is a set of nodes (i.e. genes), and *E* is a set of edges (i.e. interactions between genes). In this study we used three signalling and metabolic pathway databases that are considered as references in systems biology: (i) KEGG, the Kyoto Encyclopaedia of Genes and Genomes Pathway database (34); (ii) NCI, the National Cancer Institute / Nature Pathway Interaction Database (35); and (iii) Reactome (36,37). We then used the R/Bioconductor *graphite* package to convert biological pathways into graphs of interacting genes (see (38) for more details on this procedure).

### Computation of summary statistics on gene networks

To characterise the structure of networks and check for potential differences between databases, we generated the distributions of three standard summary statistics for each of the three databases. We thus computed for each network i) the number of nodes, ii) the number of edges, and iii) the graph density. The graph density *d* is a measure of connectivity between the nodes of the network, and it is defined as the number of edges in a set *E* compared to the maximum number of possible edges between nodes in a set *V*, therefore *d* = 2 * |*E*| / (|*V*| * (|*V*| − 1)), where |*X*| represents the number of members of a set *X*. We also analysed the overlap between pathway databases by computing the number of genes they share. Finally, we quantified the redundancy between pathways within a database by computing Jaccard’s similarity index. For a pair of networks A and B with sets of nodes *V*_A_ and *V*_B_, Jaccard’s index is defined as *J*_AB_ = (|*V*_A_| ∩ |*V*_B_|) / (|*V*_A_| ∪ |*V*_B_|).

### Workflow to detect outlier subnetworks

As the detection of outlier subnetworks includes several distinct steps, we describe here our analysis pipeline. The goal of our approach is to search within each gene network the subnetwork with the largest signal of interest (e.g. evidence of selection) using a simulated annealing approach (33). Our algorithm is globally similar to that used by Ideker et al. (29), but our method differs in two important ways, as described below. First, whereas Ideker et al.’s method aimed at finding the highest-scoring *subset* of nodes, we aim here at finding the highest-scoring *subnetwork* (i.e. a subset of genes that are directly connected by edges). Second, we consider a statistical testing procedure that explicitly considers the optimization procedure when computing the p-value of a given subnetwork. Indeed, the score of a given subnetwork identified by the simulated annealing algorithm cannot be compared to that of a random subnetwork, as simulated annealing would identify a high scoring subnetwork even in absence of any true signal (39).

### Gene and subnetwork scores

We begin our testing procedure by assigning a score to each of the gene (node) in our network. In population genetics applications looking for subsets of selected genes, this score might be a measure of population differentiation between populations (e.g. *F*_ST_), the result of a selection test, or the difference in some measure between cases and controls. If this score is available for different SNPs in a given gene, we need to summarize their scores in some way, as our method assumes that each gene has a single score. For instance, the SNP with maximum score can be selected to represent a gene, or the average of the SNP-specific scores can be computed over all SNPs assigned to a gene. We then use an aggregate score for a subnetwork of size *k* following (29) as *s* = Σ(*X*_*i*_) / √(*k*), where *X*_*i*_ is the score of the *i*-th node (gene).

### Subnetwork score normalization

We then normalize the scores of subnetworks such as to be able to compare subnetworks of different sizes. Indeed, we expect to observe less variance in subnetwork aggregate scores in large than in small networks. The score of a given subnetwork of size *k* is thus normalized as *z’*_*k*_ = (*s_k_* - *μ*_*k*_) / *σ*_*k*_ where *μ*_*k*_ and *σ*_*k*_ are the mean and standard deviation of the score of a subnetwork of size *k*, computed empirically over 10,000 random subnetworks of size *k*, obtained for each data base separately. The means and standard deviations of subnetworks of sizes *k*_min_ to *k*_max_ are computed once and stored in a lookup table. Random subnetworks of size *k* are obtained by i) randomly selecting a network in the database with a probability depending on the network size, ii) randomly selecting a gene from this network as an initial member of the subnetwork, and iii) iteratively adding *k*-1 other randomly chosen genes that are connected to the growing subnetwork.

### Searching for optimal subnetworks with simulated annealing

We have used a simulated annealing algorithm to detect the Highest Scoring Subnetwork (HSS) of each gene network. The general idea is to start with a random subnetwork, and modify it progressively by adding or removing one node at a time until we reach a subnetwork with the highest possible normalized score. The algorithm takes as initial parameters the number of iterations *N* to perform and the annealing parameter alpha, which determines a temperature function *T*(*α*) that decreases geometrically over time.

In more details, our search algorithm is as follows:

1. Select a starting subnetwork of arbitrary size *k*_min_, defined at random.
2. Calculate its normalized score *z*_*i*_.
3. Modify the current subnetwork: First, select a node at random from the following list: i) nodes not belonging to the current subnetwork, but that are connected to it by a single edge, ii) terminal nodes of the current subnetwork, iii) internal nodes of the current subnetwork which are not articulation points (i.e. whose removal will not create two disjoint subnetworks). If the selected node is not part of the current subnetwork then add it, else remove it.
4. Calculate the new subnetwork's normalized score *z*_*i*+1_
5. Accept the new subnetwork with probability min(1,*p*), where the annealing probability *p* = exp([*z*_*i*+1_ - *z*_i_] / *T*_*i*+1_), and *T*_*i*+1_ is the annealing temperature for the iteration *i*+1. This typical simulated annealing equation means that a new subnetwork is always accepted if its normalized score is larger than that of the previous subnetwork, and that less optimal subnetworks are more and more difficult to accept with more iterations of the algorithm.
6. Repeat steps 3-5 above for a given predefined number of iterations *N*.
7. Record the resulting subnetwork and its score.

This algorithm is expected to find the global optimum for a sufficient number of iterations (29), but as its performance could vary between networks, we have run it five times for each pathway, and recorded the subnetwork with the highest score.

### Statistical testing procedure

To test if the score of the estimated HSS is significantly larger than what would be expected by chance, we need to generate the null distribution of HSS for subnetworks of a given size. To do this, we cannot simply randomly sample subnetworks and compute their scores in the original dataset, as we need to take into account the fact that the optimization procedure will bias the subnetwork scores towards high value. To take this effect into account, we have generated a null distribution of optimised scores. To do this, we first permute gene scores across all networks of a given database. Then a network is randomly chosen with a probability proportional to its size, and the optimisation algorithm is applied to obtain the HSS on the permuted dataset. The score of the resulting HSS is finally recorded. This procedure is repeated *N* times to generate the null distribution. The empirical p-value of a given observed HSS is then obtained as the proportion of random HSS of similar size that have a score larger or equal to the observed HSS score (unilateral test).

As many subnetworks are tested with our procedure, we have corrected the inferred p-values for multiple testing by computing q-values, which are false discovery rates (FDRs) that would be computed if the observed p-value was used as a threshold to declare significance. To do this, we used the FDR method (40) implemented in the R package *qvalue*.

### Pipeline implementation

Our analysis pipeline has been implemented in R, and graphical representations of the networks and HSS were made using the software Cytoscape (41,42), called from R with the Bioconductor package RCytoscape (43).

### Test of the method on simulated data

As simulated annealing is an approximate method, we studied its performance using a simulation-based approach. We simulated pseudo-observed data sets by building a random network of size *N* using a random edge model, i.e. where an edge is drawn with a given probability *p*. Then, a connected subnetwork of size *k* is randomly sampled within the network. The node scores from the subnetwork are drawn from a normal distribution *N*(*μ*_HSS_,1), where *μ*_HSS_ is the average score of this subnetwork. The score of the other nodes of the network are drawn from a standard normal distribution *N*(0,1). We then apply our simulated annealing algorithm to find the highest-scoring subnetwork using with *i* iterations. Therefore, the outcome of our search depends on five parameters: the network size *N*, the HSS size *k* and its average expected score *μ*_HSS_, the network connectivity *p* and the number of iterations *i*.

In order to characterize the accuracy of our network search and to better understand which parameters have an impact on our estimation, we computed, for each simulation, the number of true positives (*TP*, the number of nodes from the true HSS that are correctly identified), the number of true negatives (*TN*, the number of nodes that are not in the HSS and that are not identified), false positives (*FP*, the number of nodes wrongly identified as part of the true HSS) and false negatives (*FN*, the number of nodes from the true HSS that have not been identified). We then computed two measures of performance: precision or positive predictive value: *PPV* = *TP* / (*TP* + *FP*); and sensitivity or true positive rate: *TPR* = *TP* / (*TP* + *FN*).

To assess the impact of our five parameters on the precision (*PPV*) and on the sensitivity (*TPR*) of our estimation, we used a Generalised Linear Model (GLM) where the response variables are the counts of *TP* and *FP* for precision, and the counts of *TP* and *FN* for sensitivity. The predictor variables are the five above-mentioned parameters, and the error follows a binomial distribution.

To test the performance of our significance testing procedure that explicitly takes the optimisation process into account, we computed p-values using the null distribution obtained from 10,000 runs of simulated annealing on data generated with *μ*_HSS_ = 0 (i.e. the null hypothesis).

### Application to real data: detection of convergent adaptation to altitude in humans

We analysed a dataset published by Foll et al. (24) on convergent adaptation to altitude in Tibetans and Andeans. This data set consists of 632,344 SNPs genotyped in four populations: two populations living at high altitude in the Andes (49 individuals) and in Tibet (49 individuals), as well as two lowland related populations from Central America (39 Mesoamericans) and East Asia (90 individuals). For each SNP, a probability of convergent adaptation has been computed under a hierarchical Bayesian model (24). To get a unique score per gene, as required in our methodology, we used the p-value of the highest scoring SNP mapped within a gene or less than 50 kb away.

We applied our methodology to detect subnetworks under selection on this dataset. The three pathway databases were analyzed separately (i.e. every step of the workflow has been done independently for each database). Pathways for which the largest connected subnetwork size was less than *k*_min_ = 5 nodes were removed from the analysis, since we wanted to avoid focusing on small subnetworks. Aggregate subnetwork score distributions have been generated by sampling 10,000 random subnetworks for each possible size *k*. The HSS search algorithm has been applied to every pathway with *N* = 10,000 simulated annealing iterations. The p-value of the obtained HSSs was inferred from the distribution of scores of 10,000 random HSS generated under the null hypothesis (i.e. permuted data).

## RESULTS

### Test of the method on simulated data

We first studied the performance of our approach in terms of precision (i.e. the fraction of selected genes in the estimated highest-scoring subnetworks (HSS)) and sensitivity (i.e. the proportion of selected genes that are identified as such) by analysing pseudo-observed data. We generated random networks and HSS based on five parameters (see Material and Methods). We ran our algorithm on the simulated data and compared the estimated HSS to the true HSS. Using logistic regressions, we show that out of the 5 parameters tested, 4 have a significant impact on both the precision and sensitivity of the method (Table 1). Most of the model deviance is explained by the mean score of the selected genes, the network size, and the subnetwork size. As expected, precision goes up with *μ*_HSS_ and the false positive rate is lower than 0.05 when *μ*_HSS_ > 4 (Figure 1A). Furthermore, even if network (*N*) and subnetwork (*k*) sizes influence our ability to correctly identify HSS, *N* and *k* have a negligible impact on the precision of our estimations when the true subnetwork score is sufficiently large. Indeed, in this case precision remains high for a broad range of *N* and *k* values (Figure 1B and 1C). Even though one would have thought that the number of iterations of the simulated annealing algorithm was an important parameter for the success of the algorithm, it has a limited impact (Table 1) and 5,000 iterations appear enough to achieve high precision (Table 1). Finally, we find that network density has no real influence on the performance of our method.

**Figure 1:**
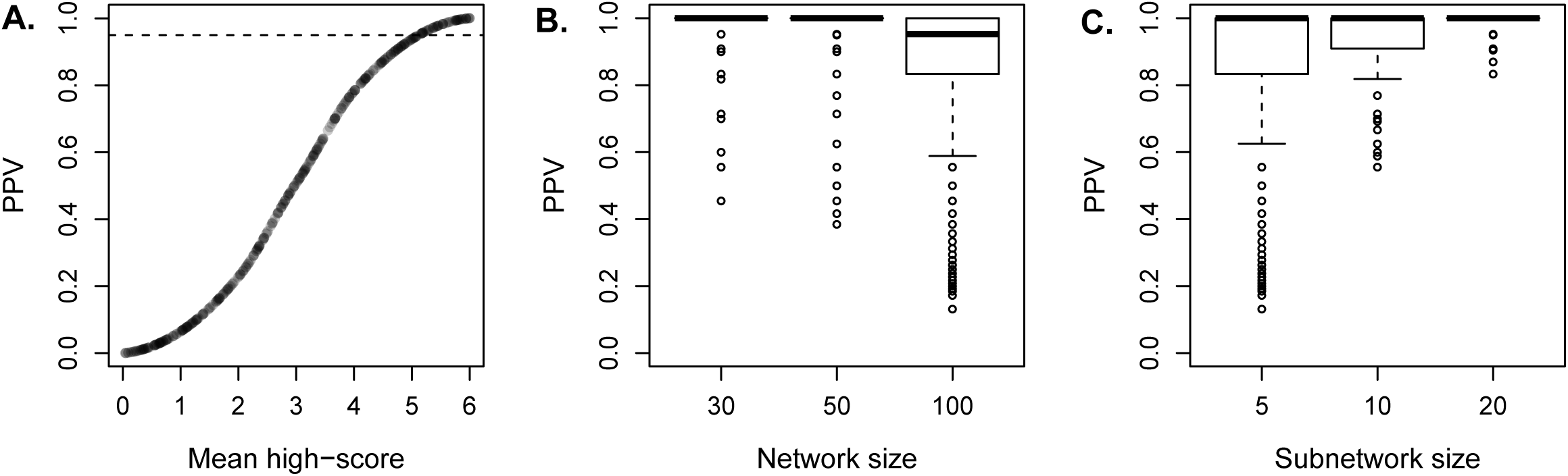
Impact of different parameters on the precision of the estimation (PPV). The predictions of the GLM for the influence of *μ*_HSS_ on the precision is represented (A), as well as the PPV as a function of network size (B) and subnetwork size (C) when *μ*_HSS_ is fixed to 5.

**Table 1:** Estimates of the effects of the five parameters on the precision (PPV) and sensitivity (TPR) obtained under a logistic regression framework. For each parameter, the coefficient, p-value and the percentage of explained total deviance (%TD) are indicated.

|  | PPV |  |  | TPR |  |  |
| --- | --- | --- | --- | --- | --- | --- |
|  | Estimate | p-value | %TD | Estimate | p-value | %TD |
| $N^1$ | -0.025 | $< 2.10^{-16}$ | 17.5 | $4.3.10^{-3}$ | $< 2.10^{-16}$ | $< 1$ |
| $k^2$ | 0.14 | $< 2.10^{-16}$ | 21.3 | -0.03 | $1.4.10^{-15}$ | $< 1$ |
| $\mu_{HSS}^3$ | 0.75 | $< 2.10^{-16}$ | 45.8 | 1.29 | $< 2.10^{-16}$ | 66 |
| $d^4$ | -0.029 | 0.29 | $< 1$ | $5.10^{-2}$ | 0.31 | $< 1$ |
| $i^5$ | $-1.8.10^{-6}$ | $2.10^{-5}$ | $< 1$ | $1.5.10^{-6}$ | 0.05 | $< 1$ |

<sup>1</sup>Network size; <sup>2</sup>HSS size; <sup>3</sup>HSS mean score; <sup>4</sup>Network density; <sup>5</sup>Number of iterations

Then, in order to verify that our statistical testing procedure behaves properly, we computed the p-value distribution under the null hypothesis of *μ*_HSS_ = 0. In that case, p-values do not depart significantly from a uniform distribution (Kolmogorov-Smirnov test, *D* = 0.03, *p* = 0.76; Figure S1), which is the behaviour expected when the null hypothesis is true.

### Pathways databases characteristics

We used pathways defined in three databases: KEGG, NCI and Reactome (including respectively 225, 189 and 1095 pathways). To see whether we should treat these databases separately or not, we first computed statistics summarizing the main properties of these databases. First, we characterized the overlap between these databases, i.e. the number of genes shared between databases. We show that even if they substantially overlap in their gene content, the three databases have a large number of private genes (Figure S2A). We also characterized the overlap between pathways within databases using Jaccard’s index. We computed the redundancy within a database as the proportion of pathways pairs with an overlap higher than a given threshold as a function of this threshold (44). We find that pathways from the three databases have different levels of overlap, with Reactome having the largest fraction on non-overlapping pathways (Figure S2B). Finally, we computed summary statistics to understand the structures of the networks in the different databases. The distributions of the number of nodes, the number of edges and the connectivity are also strikingly different between the three databases (Figure S3). Since pathways of these three databases had different properties, we have analysed them separately, using genes from each database to build separate null distributions and perform statistical tests.

### Adaptation to altitude in humans

We analysed the data from Foll et al. (24), who studied adaptation in two human populations living at high altitude. For each SNP, they computed the probability of convergent adaptation to altitude in Andeans and Tibetans under a hierarchical Bayesian model, and we used this probability as our score. We define the gene score as the highest-scoring SNP within the gene or in a 50 kb surrounding window. The distribution of gene scores appears slightly different between databases (Figure S4), again justifying the separate analysis of the three databases.

To search for high-scoring subnetworks in each pathway, we first generated the aggregate subnetwork score distributions for each database and for all possible subnetwork sizes. We then searched for the high-scoring subnetwork in each pathway using 10,000 simulated annealing iterations, and we assessed their significance from a null distribution of HSSs based on 10,000 permutated data sets (see Material and Methods). Interestingly, we find that subnetwork scores tend to be lower in denser pathways. Indeed, the estimated subnetwork score significantly decreases with the density of a pathway (linear regression, *F*(1,1339) = 42.11, *p* = 1.2.10^-10^; *R* = 0.17; Figure S4). This result is unlikely to be an artefact, as our simulation study shows that our procedure is not affected by network density (Table 1). Therefore, genes potentially involved in adaptive processes seem to be preferentially found in pathways with less gene-gene interactions. These results are in agreement with other empirical studies that showed that deleterious mutations tend to accumulate at the periphery of gene networks (45). Even though positive selection can also act on genes with more interactions (31,46), this result suggests that adaptation to altitude has mainly targeted genes with less pleiotropic effects since the number of interactions of a gene is clearly correlated to its pleiotropy level (47).

We then considered a HSS as significant if it showed a p-value < 0.01 and a q-value < 0.20. None of the pathways tested in the Reactome database remained significant after multiple test correction. We identified four pathways with a significant HSS in the NCI database and six such subnetworks in KEGG (Table S1). The overall top-scoring pathway is the HIF-2-α transcription network (Figure 2), a pathway containing genes known to respond to hypoxia conditions. EPAS1 (HIF-2-α) is the top-scoring gene, it is a transcription factor active under hypoxic conditions. All the other significant genes within this pathway are directly interacting with EPAS1 and should thus play an important role in response to hypoxia. Some of these genes are inhibitors (CITED2) or cofactors (ARNT) of Hypoxia-Inducible Factors (HIF), others are regulated by HIF, such as VEGFA, a growth factor involved in angiogenesis.

**Figure 2:**
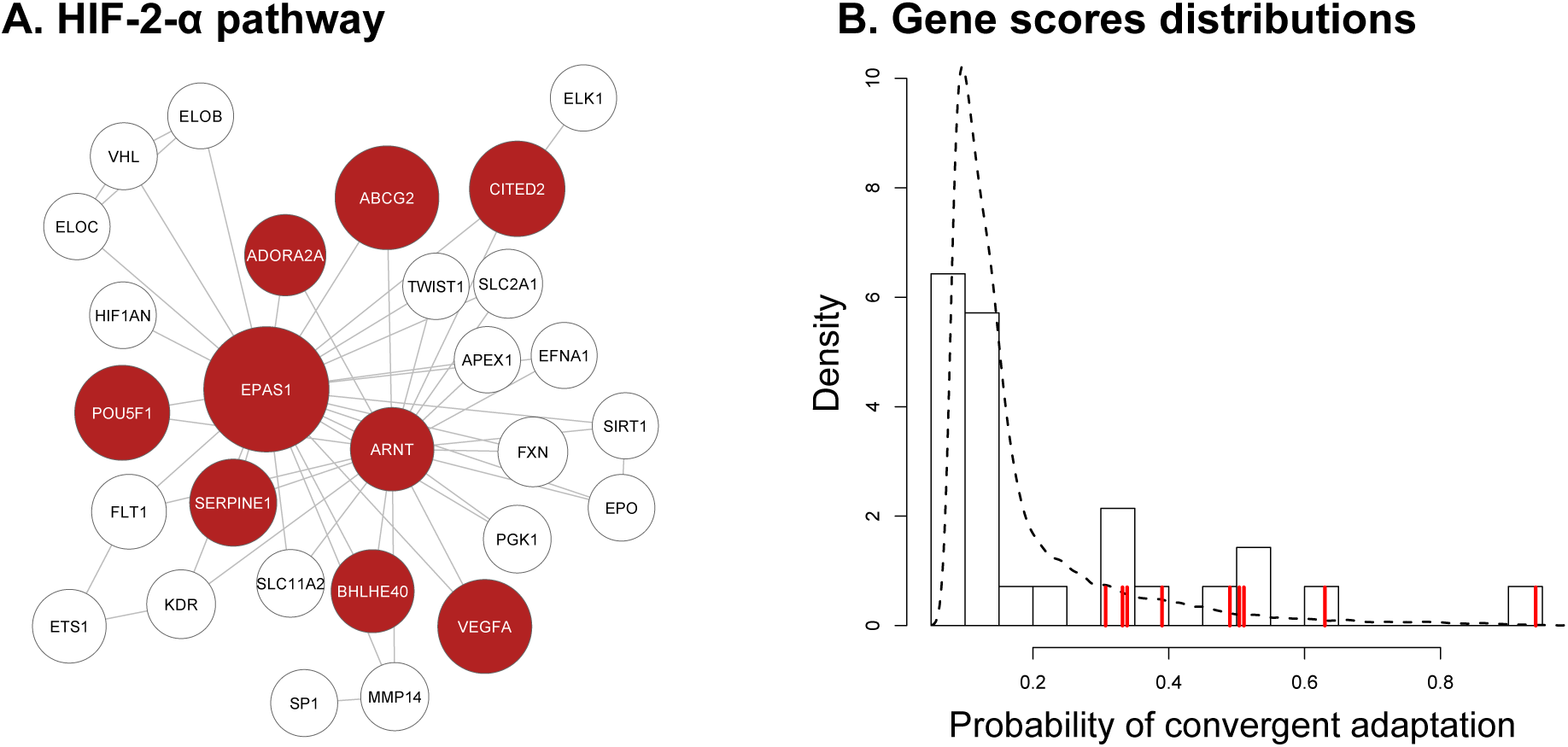
Most significant subnetwork among the three pathway databases. The HIF-2-α transcription pathway is represented as a graph (A), where each node is a gene, and the node size is proportional to the gene score. The highest scoring subnetwork (HSS) of the pathway is shown in red. The gene scores density distribution in this pathway is shown in (B). The dashed line represents the density of gene scores within all the KEGG database, the histogram shows the distribution of genes scores within this pathway, and the vertical red lines indicate the scores of the genes belonging to the HSS.

When top-scoring HSSs were overlapping by one or more gene, we merged them in a single network (Figure 3). After this procedure, we observe four distinct clusters of genes. First, in the NCI database, we find a single cluster of genes within four pathways involved in vascular processes such as angiogenesis, response to hypoxia or blood coagulation (Figure 3A). Among these, the top-scoring genes are Endothelial PAS domain-containing protein 1 (EPAS1), Interleukin-6 (IL6), Angiopoietin 1 (ANGPT1), Pleiotrophin (PTN), Tyrosine-protein phosphatase non-receptor type 11 (PTPN11) and Epidermal Growth Factor (EGFR). We also observe many genes in these HSS that present lower scores. Most of these are growth factors, such as genes in the Insulin Growth Factor (IGF), receptor tyrosine kinases (ErbB, EGFR), Neurotrophic Factors (NTF) or Interleukin (IL) families. We identified three other clusters of genes in the KEGG database that are involved in very different biological processes (Figure 3B). First, a large network of 32 genes involved in metabolic functions where the top-scoring genes are Alcohol Dehydrogenase (ADH) subunits, most of the other genes being other aerobic metabolism related enzymes such as the Glucuronosyltransferase (UGT), Glutathion S-tranferase (GST) or Glutamic-Oxaloacetic Transaminase (GOT) families. All of them present moderate probabilities of convergent adaptation (< 0.8). Second, an immunity-related cluster is observed, including 6 Human Leucocyte Antigen (HLA) genes. A last cluster consists in three genes related to neuronal cell growth, with Neuroligin 4 (NLGN4X) being the top-scoring gene in this database.

**Figure 3:**
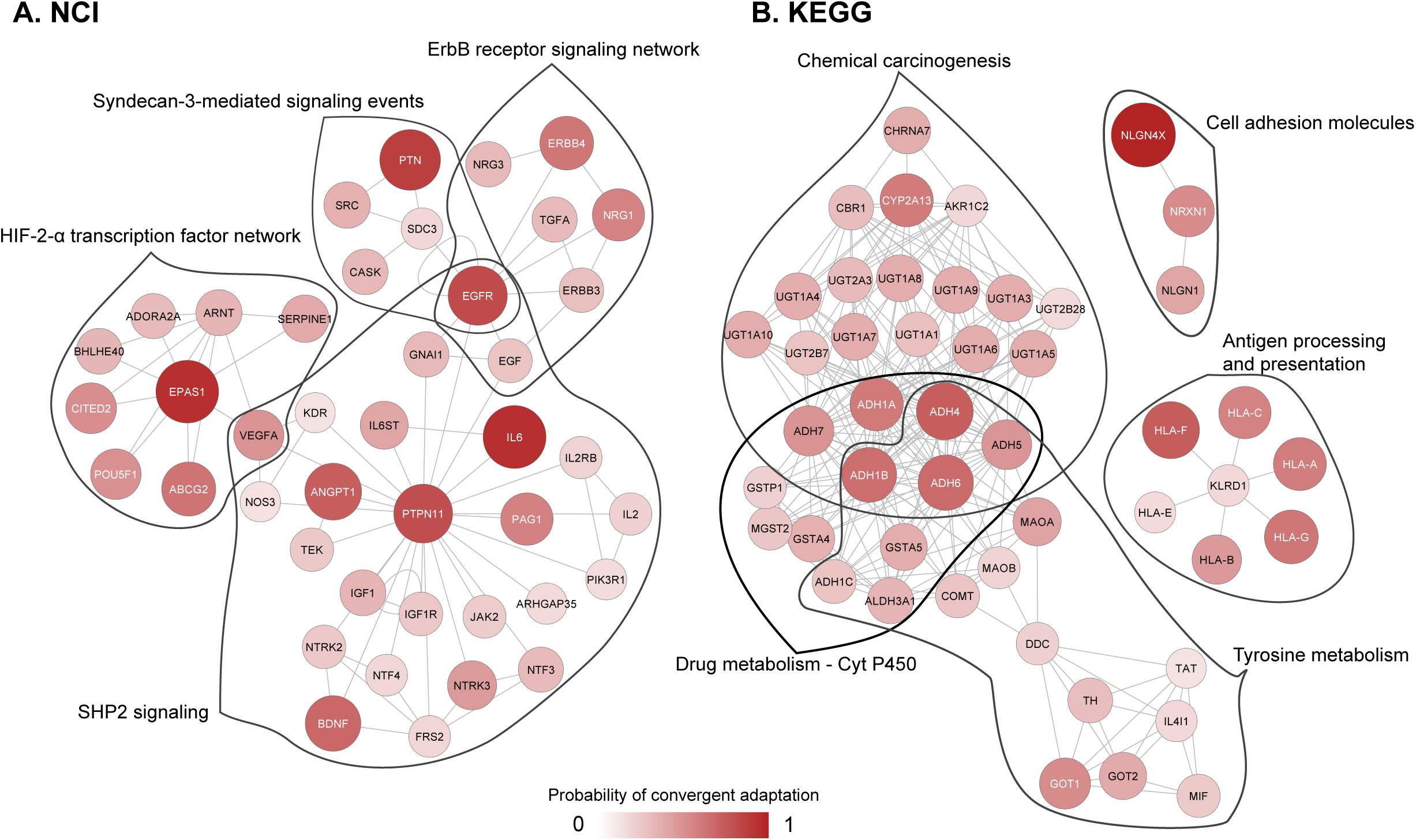
Merged significant subnetworks. For each database, NCI (A) and KEGG (B), we merged the significant subnetworks of genes if they overlapped. The colour intensity and size of the nodes are proportional to the gene score. Red lines delimit the individual significant subnetwork and the names of pathways to which they belong are shown next to it.

## DISCUSSION

### New insights into human adaptation to altitude

The challenges of living at high altitude impose a very strong selective pressure on individuals, mainly due to low oxygen levels leading to hypoxia (48). Physiological changes have been identified in Tibetans and Andeans living at high altitude (49), and many studies have unveiled the genetic bases of these physiological changes (reviewed in (48)). Adaptation to altitude thus offered us a good positive control to test our new method on real data, and therefore, the fact that our top subnetwork is found in the HIF-2-α transcription pathway is reassuring. This pathway is indeed a key component of the response to hypoxia, as it modulates or induces various physiological responses such as angiogenesis, haemoglobin concentration or erythropoiesis (50). Numerous genes within this pathway have already been proposed to be under selection in Tibetans and Andeans, such as EPAS1 and IL6 (50-53). In addition to these usual suspects, we identify many other genes with scores that remain below the detection threshold of the original genome scan (24), and which show a much more moderate signal of convergent adaptation. The identification of other candidate genes present in the HIF pathway is in line with the view that adaptation to altitude has a polygenic basis (50). For instance, we identified pleiotrophin (PTN), which acts as an angiogenic factor through multiple mechanisms (54), but which has to our knowledge never been identified as a major player in adaptation to altitude. Another gene, PTPN11, has a high score and a central position in one of the significant subnetworks. It encodes the protein tyrosine phosphatase SHP-2, which regulates heart and blood cells development during embryogenesis, as well as other tissues (55). The Cell Adhesion Molecules (CAM) pathway also presents interesting signals, as we have identified a small cluster of 3 genes coding for neuroligins, which are neuronal proteins involved in the modulation of synaptic transmission (56). However, it has recently been shown that these genes are also involved in vascular processes (57,58). The NLGN1 gene thus seems to be a strong candidate for adaptation to high-altitude in Tibetans and Andeans and mechanisms linked to neuroligins action in angiogenesis at high altitude would deserve further investigation.

Note that we have also identified a cluster of genes involved in separate metabolic processes. Signals of adaptation at ADH and ALDH genes have been observed in the original study as well as in Ethiopian populations living at high altitude (59). As suggested in the original study, these genes could be involved in fatty-acid degradation and energy production in the mitochondrion: in case of hypoxia, alternative pathways such as omega-oxidation (including ADH genes) could be an alternative to beta-oxidation (24).

### Advantages and limitations of the method

The search for high-scoring subnetworks is a combinatorial optimization problem for which several methods have been developed (29,39,60). Here, we describe a new method to detect selection in biological pathways based on a simulated annealing algorithm that extends a previous approach (29) by searching for the highest scoring *subnetwork* of interacting genes rather than for the highest scoring subset of nodes, i.e. we constrain the search to a single connected set of genes. Even though an exact algorithm has been developed to find the optimal subnetwork, it is not generally applicable, as it can only be applied to a list of p-values coming from a mixture of beta distribution (39). On the other hand, our simulated annealing method does not require any assumption on the distribution of gene scores, and it can therefore be applied to a wider range of problems. In addition, our statistical testing procedure explicitly takes into account the optimization procedure, by building a null distribution of high-scoring subnetworks in permuted data. The generation of this null distribution is a crucial step to prevent simulated annealing to identify subnetworks in the absence of any signal (39), and we show here by simulation that our statistical procedure behaves properly in terms of type I error (Figure 1 and S1).

An interesting feature of our approach is the integration of functional information into the analysis by directly testing biologically relevant gene sets. This procedure allows one to better interpret the output of a genome scan and to find the potential functions that are involved in the adaptive process. This is in clear contrast with traditional Gene Ontology (GO) analyses (61) that are typically performed on a list of top scoring candidate genes. For instance, a GO analysis performed in the original study (24) reveals only 2 significant GO terms: ‘‘ethanol oxidation’’ (GO:0006069) and ‘‘positive regulation of transmission of nerve impulse’’ (GO:0051971). This GO analysis thus missed some important biological processes involved in adaptation to altitude. Note that 14 of the 72 genes initially identified as candidates for adaptation to altitude are also present in our significant HSSs (Figure 3). 46 of the 72 top scoring genes are not included in our current analysis, either because they are absent from the pathways databases (n = 31), or because no SNP could be associated to them (e.g. because the closest SNP is more than 50 Kb away from them) (n = 15). Therefore, only 26 genes were identifiable with our method. The fact that a large fraction (46%) of these genes are not present in our significant HSSs and that only 4 out of 9 HSSs include a top-scoring gene shows that our method is not just agglomerating less significant genes around top scoring genes. In any case, our results seem biologically more relevant than a GO analysis output and in this case easier to interpret.

Our approach is conceptually close to the method developed by Daub et al. (7), which consisted in testing if a whole pathway presented a shift in the gene score distribution. The main difference with this previous approach is that we aim here at finding high-scoring subnetworks within pathways. Indeed, it is more likely that polygenic adaptive events have focused on only a subset of genes rather than on a whole pathway. In addition to be able to identify small subsets of genes even in large pathways, our approach allows one to identify outlier functions and genes at the same time, whereas under the previous whole-pathway approach, pathways had to be manually inspected in order to know which genes were driving adaptation (7). However, Daub et al.’s approach (7) has some advantages as it can be applied to any pathway, as it is not limited to pathways for which gene interaction networks are explicitly available. Therefore, the two approaches should be seen as complementary.

Whereas the present methodology overcomes some common problems associated to genome scans for selection, such as being able to identify genes with moderate selection score (Figure 2 and 3), and to explicitly associate candidate gene-sets to biological functions, it also presents some limitations as compared to other methods to detect selection. For instance, our approach is limited by the availability of pathway and network information. Therefore, some genes and biological functions cannot be tested, and the method is not easily applicable to non-model species for which no pathways databases are available. Then, one should be aware that only a certain type of biological functions are tested, i.e. biochemical phenotypes, and we thus have no information about higher-order phenotypes, e.g. height or weight. Finally, this method does not allow one to identify isolated top-scoring genes. However, such isolated outlier genes are easily identified with a classical genome scan. One can thus check if outliers are represented among significant subnetworks and therefore determine if selection has only targeted these single genes of if higher-order processes have been the target of selection.

Overall, our method allowed us to study an example of human adaptation from a gene network perspective. Based on information about gene interactions and a proxy for selection, we were able to identify potential undiscovered targets of selection, like pleiotrophin or neuroligins. This method has thus the potential to detect new genetic bases of adaptation in humans, as well as in other species for which gene interactions databases exist or could be inferred. In addition, even though we have applied this search algorithm to a case of human evolution, the same workflow can be used in other fields, such as for the study of differential gene expression, GWAS or any kind of analysis for which a score can be obtained for any given gene.

## AVAILABILITY

Our approach has been implemented as a fully automated R package. The source code and documentation are available on Github (http://www.github.com/CMPG/signet).

## SUPPLEMENTARY DATA

Supplementary Data are available at NAR online.

## ACKNOWLEDGEMENT

We would like to thank Isabelle Dupanloup for bioinformatics support.

